# The kinetics of nucleotide binding to isolated *Chlamydomonas* axonemes using UV-TIRF microscopy

**DOI:** 10.1101/547091

**Authors:** M. Feofilova, M. Mahamdeh, J. Howard

## Abstract

Cilia and flagella are long, slender organelles found in many eukaryotic cells where they have sensory, developmental and motile functions. All cilia and flagella contain a microtubule-based structure called the axoneme. In motile cilia and flagella, which drive cell locomotion and fluid transport, the axoneme contains, along most of its length, motor proteins from the axonemal dynein family. These motor proteins drive motility by using energy derived from the hydrolysis of ATP to generate a bending wave, which travels down the axoneme. As a first step towards visualizing the ATPase activity of the axonemal dyneins during bending, we have investigated the kinetics of nucleotide binding to axonemes. Using a specially built UV-TIRF microscope, we found that the fluorescent ATP analog mantATP (methylanthraniloyl adenosine triphosphate), which has been shown to support axonemal motility, binds all along isolated, immobilized axonemes. By studying the recovery of fluorescence after photobleaching, we found that there are three mantATP binding sites: one that bleaches rapidly (time constant ≈ 1.7 s) and recovers slowly (time constant ≈ 44 s), one that bleaches with the same time constant but does not recover and one that does not bleach. By reducing the dynein content in the axoneme using mutants and salt extraction, we provide evidence that the slow-recovering component corresponds to axonemal dyneins. The recovery rate of this component, however, is too slow to be consistent with the activation of beating observed at higher mantATP concentrations; this indicates that the dyneins may be inhibited due to their immobilization at the surface. The development of this method is a first step towards direct observation of the travelling wave of dynein activity.

## Introduction

Motile eukaryotic cilia and flagella are dynamic structures that move in a periodic wave-like pattern to propel cells through liquid and to move fluid across cell surfaces (1). At the core of cilia and flagella is the axoneme, a bundle of nine doublet microtubules surrounding a core of two single microtubules, together with hundreds of other associated proteins (2, 3). Among these associated proteins are the axonemal dyneins, which are located between the doublets along the length of the axoneme (4). The dyneins use energy derived from the hydrolysis of ATP to slide microtubule doublets (5–7) and the sliding is converted to bending by constraints that prevent sliding at the base. The dyneins undergo an ATPase cycle: ATP molecule binds, is hydrolyzed and the products (ADP and organic phosphate) are released (8, 9). The ATPase cycle is thought to be coupled to force generation through conformational changes of the dyneins (10–12). The similarity of the frequency of the axonemal bending wave to the ATPase rate of the axonemal dyneins suggests that each dynein ATP cycle is coupled to a cycle of doublet sliding and bending (13). In this way, dynein is thought to power the axonemal beat.

Dyneins are believed to generate sliding forces all along the length of the axoneme. First, the dyneins are present all along the length (14). And second, early modeling studies, which predated the discovery of dynein, argued that the bending wave was caused by a wave of activity that traveled all along the length: if motors were only active at the base, driving a whip-like motion of the axoneme, then the amplitude of the bending wave would decrease as it propagates (15), in contradiction to observations. The hypothesis that the axonemal bending wave is driven by a traveling wave of dynein activation forms the basis for many subsequent models, which can accurately predict the axonemal waveform (15–19). However, this hypothesized wave of dynein activity has never been directly measured or observed. Direct observation of a traveling wave of dynein activation is important, not just to test the models, but also to provide information about the extent of the modulation of the dynein activity, as well as its spatial and temporal properties.

One reason why it is difficult to observe directly the postulated traveling wave of dynein activation is that it is difficult to assay the activity of dynein. One experimental avenue is electron cryo-microscopy. A limitation of this approach, however, is that it is difficult to rapidly freeze a beating cilium or axoneme. For example, in a recent paper (11), an active axoneme was rapidly frozen, but during the sample preparation, which took a few seconds, it is possible that beating stopped or was altered. Another avenue is to use fluorescence to monitor the binding and unbinding of ATP. However, this approach is difficult because dyneins have a higher specificity for nucleotides than other motors such as myosins and kinesins: only a small number of ATP analogs have been shown to support force-generation by axonemal dyneins (20–22). Indeed, only the fluorescently-labeled nucleotides mantATP (2’-(or 3’)-O-(N-Methylanthraniloyl) Adenosine 5’-Triphosphate) and ant-ATP (2’- (or 3’)-O-Anthraniloyl-adenosine-5’-triphosphate) can support reactivation of isolated, demembranated axonemes, although at frequencies significantly reduced from that of ATP (21, 23). MantATP also supports translocation of microtubules in gliding assays in which outer-arm dyneins are bound to the glass surface of a perfusion chamber and microtubules introduced into the solution are observed to glide along the surface (24). While the fluorescence of mantATP has been used in solution studies on axonemal dyneins (10), mantATP fluorescence has not been observed in the intact axoneme structure. This is partly due to the low quantum efficiency of the dye and partly due to the excitation and emission spectra of mantATP, which have peaks at 355 nm and 448 nm respectively (25) and make detection difficult due to the high background signal.

Recent developments in highly sensitive cameras for use with microscopy allow imaging of low-emitting dyes such as mantATP. Here we report, for the first time, the direct observation of binding of mantATP to intact immobilized axonemes using UV Total Internal Reflection Fluorescence (TIRF) microscopy.

## Methods

### Reagents

MantATP (2’-(or-3’)-O-(N-methylanthraniloyl) adenosine 5’-triphosphate, trisodium salt) was purchased from Jena Bioscience (Jena, Germany). All other reagents were purchased from Sigma Aldrich (St. Louis, MO, USA) unless stated otherwise.

### Chlamydomonas cells

*Chlamydomonas reinhardtii* cells (CC-125 wild-type mt+ 137c) were grown in liquid tris-acetate-phosphate (TAP) medium: 20 mM tris, 7 mM NH_4_Cl, 0.40 mM MgSO_4_, 0.34 mM CaCl_2_, 2.5 mM PO^−3^_4_, and 1000-fold diluted Hutners trace elements (14). The medium was titrated to pH 7.0 with glacial acetic acid.

### Axonemes

Axonemes were purified using the method described in Alper et al (26).

*Chlamydomonas* cells were harvested by centrifugation at 900 × g for 5 min. They were then deflagellated by incubation with 4.2 mM dibucane-HCl for 90 seconds. The flagella were separated from the cell bodies by centrifugation at 24000 × g for 20 min on a 30% sucrose cushion. Flagella were then concentrated by re-suspending the pellet in 10 mL of HMDE buffer (30 mM HEPES, 5 mM MgSO4, 1 mM DTT, and 1 mM EGTA, titrated to pH 7.4 with KOH) with addition of 0.4 mM Pefabloc. The flagella were then demembranated by adding 0.2% IGEPAL CA-630 and washed in HMDE buffer. Salt-extraction of axonemes was performed by incubating with the indicated amount of KCl (0.3M - 1M) + HMDE for 20 minutes; the extracted axonemes were then centrifuged at 31000 × g for 20 minutes to remove the salt and extracted protein. The axonemes were re-suspended in HMDE buffer.

### Measurement of dynein content

Serial dilutions of each kind of axoneme were loaded on SDS-PAGE gels, 4%–15% precast Mini-PROTEAN TGX Gels, tris/tricine/SDS running buffer, and SDS sample buffer (BioRad, Hercules, CA). Gels were then stained with Coomassie brilliant blue (Life Technologies, Carlsbad, California, United States). Images of gels were obtained by scanning the gel on an Epson V700 document scanner (Epson, Suwa, Nagano, Japan). The gels were analyzed in Fiji image analysis software (16). The amount of dynein and tubulin was measured by integrating the density of the respective bands in the scanned gels. The dynein content was defined as the dynein signal divided by the tubulin signal, the latter serving as a loading control.

### Imaging

Imaging was performed using a Zeiss Axiovert 200M (Zeiss, Germany) microscope with a Zeiss alpha Plan-Fluar 100x/1.45 oil objective and a home-built laser-TIRF line. An ultraviolet diode-pumped laser (model Zouk 05-01 355nm, max power 10 mW, (Cobolt, Sweden) was used as the light-source. Images were recorded with an iXon 887 EMCCD back-illuminated camera.

For imaging, axonemes in HMDEKP buffer (30 mM HEPES, 5 mM MgSO_4_, 1 mM DTT, and 1 mM EGTA, 50 mM K-acetate, 1% w/v PEG titrated to pH 7.4 with KOH) were infused into an experimental chamber of depth 0.1 mm formed by a coverslip and a microscope slide; the glass surfaces we easy-cleaned (27).

During the bleaching phase, the field of view was continuously exposed to light for 12 seconds. During the recovering phase, the field of view was illuminated by brief flashes of the same intensity and of duration 100 ms; the intervals between flashes varied between 5 and 25 s.

### Image analysis

Collected image sequences were analyzed in MATLAB (MathWorks) with a semi-automated procedure. Line profiles were drawn along the axonemes, and the lines were widened by performing a dilation operation with a 5-pixel square structuring element to produce an axonemal region of interest. The pixels in the original image in this region were then averaged to obtain the axonemal signal (*F*_axoneme_). Equivalent regions on both side of the axonemes were used to calculate a background signal (*F*_background_). Because the background illumination varied across the field of view, due to variation in the laser intensity, we normalized the axoneme signal by first subtracting the background and then dividing by the background signal to obtain a relative axoneme fluorescence intensity:

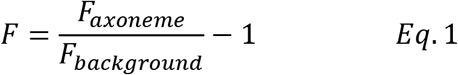

### Fitting the beaching and recovery phases

The data were analyzed separately for the bleaching phase 0 < *t* < *t_off_*, where the field of view is constantly exposed to the laser light, and the recovery phase *t* > *t*_off_, where *t*_off_ = 12 s is the time at which the continuous laser light is shuttered (Figure 1B). The bleaching phase was fitted with the empirical equation:

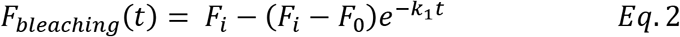

and the recovery phase was fitted with:

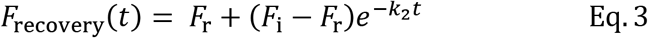

*F*_0_ is the initial relative fluorescence intensity value (at *t* = 0), *F*_i_ is the bleaching steady-state intensity, *F*_r_ is the maximum recovery intensity. *k*_1_ and *k*_2_ are the rate constants of the bleaching and recovery phases respectively. Separation into these two phases was possible because there was a large difference between the rate constants of the bleaching and recovery phases, and because bleaching had nearly reached a steady state at *t*_off_. The parameters were obtained by fitting the data to these exponential functions using non-linear least squares (MATLAB, “fit” function).

**Figure 1.**
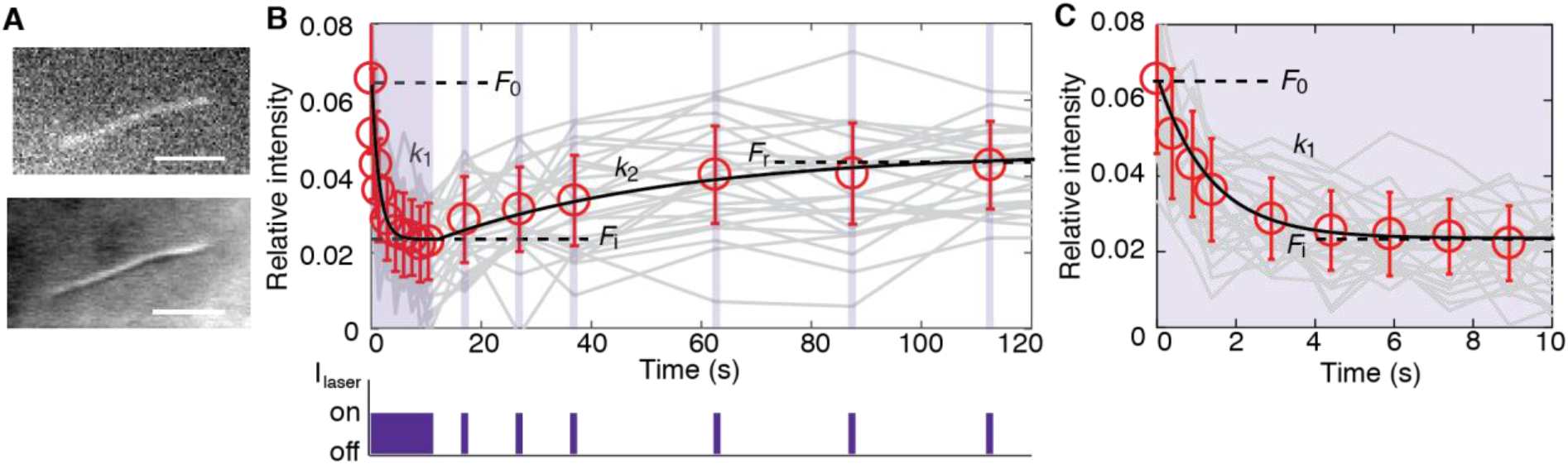
Bleaching and recovery of *oda1* axonemes under UV-TIRF. **A** First frame image of an axoneme form the experiment (top). Differential interference contrast (DIC) image of the same axoneme (bottom). Scale bars 10μm. **B** Upper panel shows a bleaching and recovery time: mean ± standard deviation shown in red, a random sampling of raw traces shown in grey. Lower panel indicates the illumination sequence: 12 second constant illumination (bleaching phase) is followed by 100-ms-duration flashes of the same intensity. C. Higher temporal resolution plot of the bleaching phase.

## Results

### MantATP binds to the axoneme

We incubated flow cells with solutions containing isolated, demembranated *Chlamydomonas* axonemes and waited a few minutes until several had bound to the surface in each field of view (82 μm × 82 μm). We then infused a solution containing 1 μM mantATP. Using a TIRF microscope with long-wavelength UV illumination (355 nm laser), we observed a fluorescent signal along the entire axoneme (Figure 1A, TIRF image in upper panel to DIC image in lower panel). The same assay performed in the absence of mantATP did not produce a fluorescent signal from the axonemes. Thus, there are mantATP binding sites all the along the axoneme, though we cannot exclude the possibility that the fluorescence is reduced within a micrometer from the ends.

To investigate the kinetics of the binding and unbinding of mantATP to the axoneme, we used a bleaching-and-recovery protocol. In this assay, the fluorescence was bleached by intense, continuous UV illumination and the recovery of fluorescence was monitored by intermittent brief light pulses. A typical trace is shown in Figure 1B in which there is partial bleaching followed by partial recovery of fluorescence. During the bleaching phase, the fluorescent signal decreased over a few seconds from its initial value *F*_0_ towards a steady-state value *F*_i_. During the recovery phase, the fluorescent signal approached the recovery value *F*_r_ over approximately one minute. See Table 1 for the bleaching and recovery rates, together with the amplitudes *F*_0_, *F*_i_ and *F*_r_. We identified two general features of the bleaching and recovery curves: (i) the bleaching rate was much higher than the recovery rate, and (ii) the recovery was always incomplete (*F*_r_ < *F*_0_).

**Table 1.** Measured parameters. Parameters measured from the bleaching and recovery curves like those shown in Figure 1 for ten separate experiments at 1 μM mantATP, with 50 - 150 axonemes analyzed in each experiment. The value of the intermediate level obtained by fitting the bleaching curves was similar to the value obtained by fitting the recovery curves (bleaching: 0.027 ± 0.003, recovery: 0.045 ± 0.003; mean ± SE, *n* = 10 experiments.

| Parameter | | Mean ± SE |
| --- | --- | --- |
| Timescales |  |  |
| Bleaching rate | $k_1$ | $0.70 \pm 0.06 \text{ s}^{-1}$ |
| Recovery rate | $k_2$ | $0.027 \pm 0.004 \text{ s}^{-1}$ |
| Relative fluorescence intensities |  |  |
| Initial | $F_0$ | $0.068 \pm 0.008$ |
| Intermediate | $F_i$ | $0.027 \pm 0.003$ |
| Recovery | $F_r$ | $0.045 \pm 0.003$ |

### The bleaching recovery data indicates that there are (at least) three nucleotide binding sites

To understand the bleaching and recovery curves, we first attempted to fit the time courses with the one-binding-site model shown in Figure 2. The idea is that mantATP* binds to a site, is hydrolyzed to mantADP* and the mantADP* unbinds at a slow rate. The (*) refers to the fluorescent nucleotide. If hydrolysis (*h*) is faster than bleaching (*b*)—justified by data on the ATPase activity of *Tetrahymena* outer-arm dynein (Holzbaur & Johnson, 1989)—then the bleaching phase corresponds to bleaching of mantADP* to mantADP†, where (†) denotes the bleached nucleotide. In this scheme, recovery is limited by the slow unbinding of mantADP†. Usually we will omit the (†).

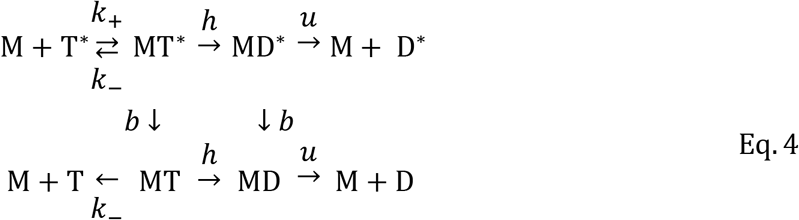

**Figure 2.** General scheme for mantATP binding, hydrolysis and bleaching. Fluorescent mantATP*, denoted by T*, binds rapidly and reversibly to a site M. mantATP* is hydrolyzed and phosphate released to produce mantADP*, denoted by D*, and both the unbleached mantADP*and the bleached mantADP†, denoted by D, unbind slowly with rate *u*, limiting the recovery rate.

This one-binding-site model cannot account for the bleaching and recovery curves shown in Figure 1 (see Appendix for the mathematical arguments). First, there should be complete recovery. Instead, the partial recovery indicates that there is a site that bleaches irreversibly. We call this the non-recovering component. Second, the slow-recovering component should exhibit almost complete bleaching because the bleaching rate (*k*_1_ = *b* + *u*, in the scheme) is much faster than the recovery rate (*k*_2_ = *u*), which corresponds to the unbinding of mantADP. Formally, 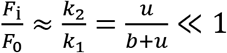(see Appendix). Instead, the bleaching is only partial (*F*_i_/*F*_0_ ≈ 0.4). This indicates that there is a site that does not bleach; we call this the non-bleaching component. Therefore, the bleaching curves indicate that there are at least three components: a slow-recovering component, a non-bleaching component and a non-recovering component.

The three-component model provides a good fit to the experimental data (Figure 3). In Figure 3B, the experimental data is shown in red circles, the fitted sum of the three components is shown in blue, and the slow-recovering, non-bleaching and non-recovering components are shown in light-blue, green and black respectively. The bleaching rate, the unbinding rate and amplitudes of the three components are given in Table 2. All three components have similar amplitudes. Because the bleaching phase did not resolve into two exponentials, the bleaching rates for the non-recovering and slow components must be similar.

**Figure 3.**
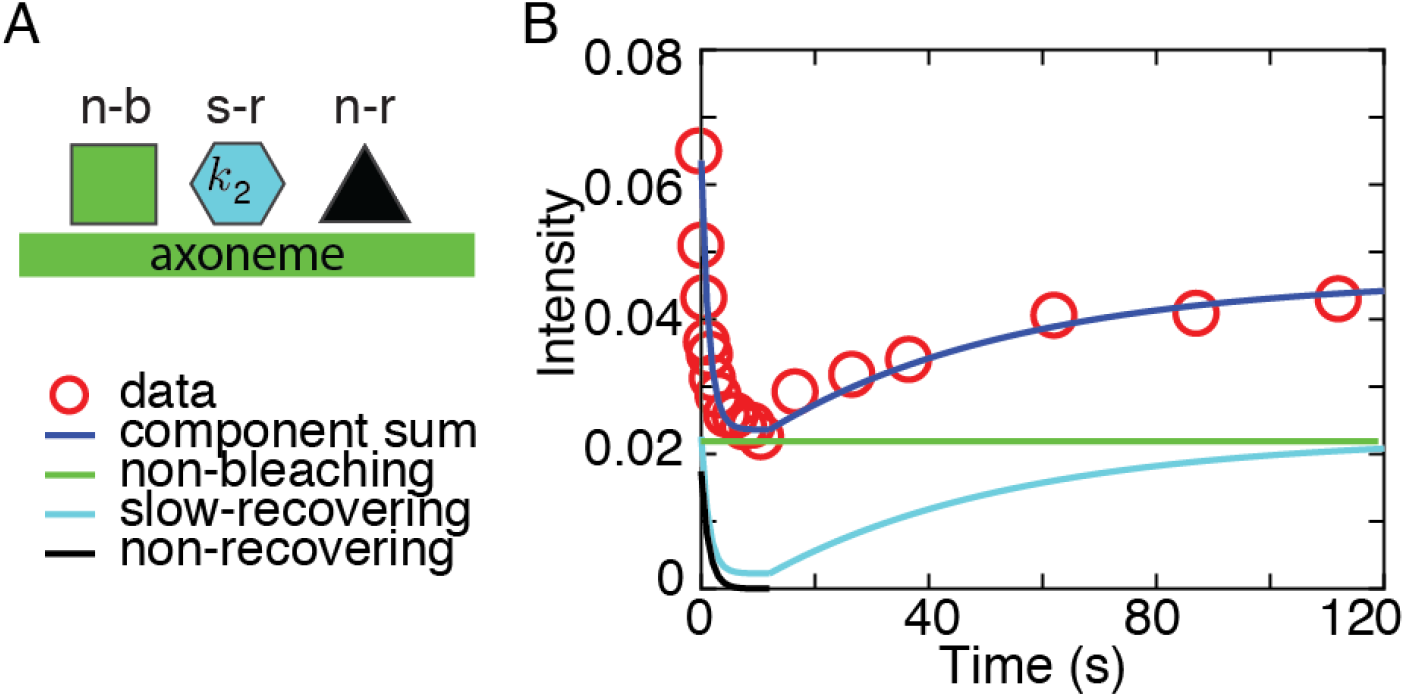
Three-component model. **A** Three binding site model with distinct kinetic characteristics. See text for details. **B** Overlay of the three-component model and experimental data. Experimental data shown in red, non-recovering component in black, non-bleaching (fixed) component in green and slow-recovering component in light blue. Sum of components is shown in dark blue.

**Table 2.** Molecular rate constants and amplitudes of the three components. The parameters from Table 1 were used to infer the properties of the three components. The non-recovering component has relative amplitude (*F*_0_ − *F*_r_)/*F*_0_ The slow recovering component has amplitude [(*F*_r_ − *F*_i_)/*F*_0_][*k*_1_/(*k*_1_ − *k*_2_)] ≈ (*F*_r_ − *F*_i_)/*F*_0_. The non-bleaching component is the rest, and is approximately equal to *F*_i_/*F*_0_.

| Parameter | | Mean ± SE |
| --- | --- | --- |
| Timescales |  |  |
| Bleaching rate constant | $b$ | $0.57 \pm 0.12 \text{ s}^{-1}$ |
| Unbinding rate constant | $u$ | $0.023 \pm 0.003 \text{ s}^{-1}$ |
| Relative amplitudes of components |  |  |
| Non-recovering | $F_x$ | $0.32 \pm 0.03$ |
| Non-bleaching | $F_f$ | $0.40 \pm 0.02$ |
| Slow-recovering | $F_s$ | $0.28 \pm 0.02$ |

### Reducing dynein in axonemes using mutants and salt extraction

To determine whether axonemal dynein is contributing to one of the three binding sites, we used axonemes with different dynein contents. One way to reduce the number of dyneins is to use mutants lacking some of the dynein arms. Another way is use axonemes whose dyneins have been extracted with KCl (14). We compared wild-type axonemes, *oda1* mutant axonemes (which are missing outer dynein arms) and salt-extracted *oda1* axonemes to further reduce the number of dyneins. The amount of dynein in each type of axoneme was measured by scanning SDS-PAGE gels. *Oda1* axonemes have less dynein than wild-type axonemes. Increasing the KCl salt concentration further decreased the amount of dynein in the axonemes (Figure 4A). Thus, by extracting the *oda1* axonemes using salt, the dynein content could be decreased by as much as 75% compared to wild-type.

**Figure 4.**
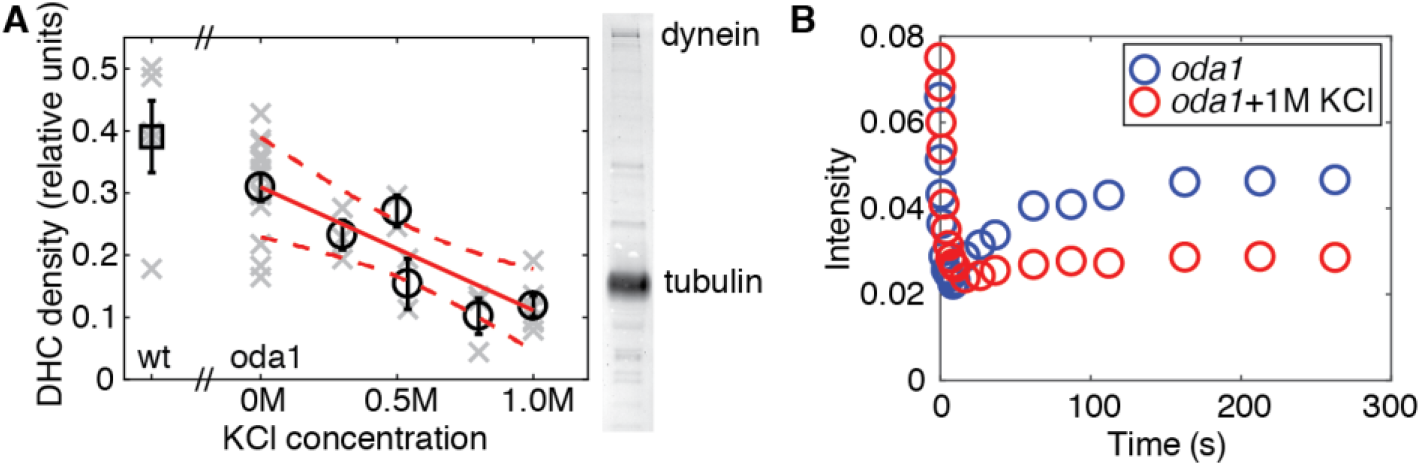
Salt extraction of dynein from the axoneme. **A** Salt-extraction of dynein from *oda1* axonemes was performed with various concentrations of KCl. Black markers represent the mean and standard error; in light gray, data from single experiments is shown, the red line is a weighted linear regression of the data (*y* = 0.31 − 0.20*x*, slope significant *t*(4) = −4.7,*p* < 0.01), dashed red line shows 95% confidence bounds. Wild-type axoneme density is represented by a square marker, the abscissa is selected arbitrarily. On the right is an example of an SDS-PAGE gel lane, showing the dynein and tubulin bands of *oda1* axoneme. **B** Example of measurements for *oda1* axonemes and *oda1* axonemes extracted with 1M KCl. A decrease in the amplitude of the recovery (slow component *F*_r_ − *F*_i_) can be seen, while the non-recovering component (*F*_i_) remains unchanged.

### Dynein reduction reduces the amplitude of the slow component

To find out whether dynein is responsible for the slow or fixed components, we compared axonemes with different amounts of dynein. In Figure 4B *oda1* axonemes and *oda1* axonemes extracted with 1M KCl are compared. The steady-state bleaching intensity *F*_i_ was similar for both kinds of axonemes while the recovery amplitude decreased significantly for the extracted axoneme. Thus, the slow component correlates with the dynein content.

To test how these findings hold across axonemes with various dynein content, we plotted the amplitudes of the non-bleaching component ≈ *F*_i_ (Figure 5A) and the slow-recovering component ≈ *F*_r_ − *F*_i_ (Figure 5B) for axonemes containing different amounts of dynein. While the amplitude of the fast component was smaller in *oda1* compared to wildtype, no further decrease occurred as the salt concentration was increased (Figure 5A *t*(6) = 1.37, *p* > 0.24). By contrast, the amplitude of the slow-recovering component decreased by more than 80%, as the salt concentration was increased and the slope was statistically significant (Figure 5B, *slope* = −0.014, *t*(6) = 8.6, *p* < .001). Thus, the slow-recovering component is likely to be axonemal dynein.

**Figure 5.**
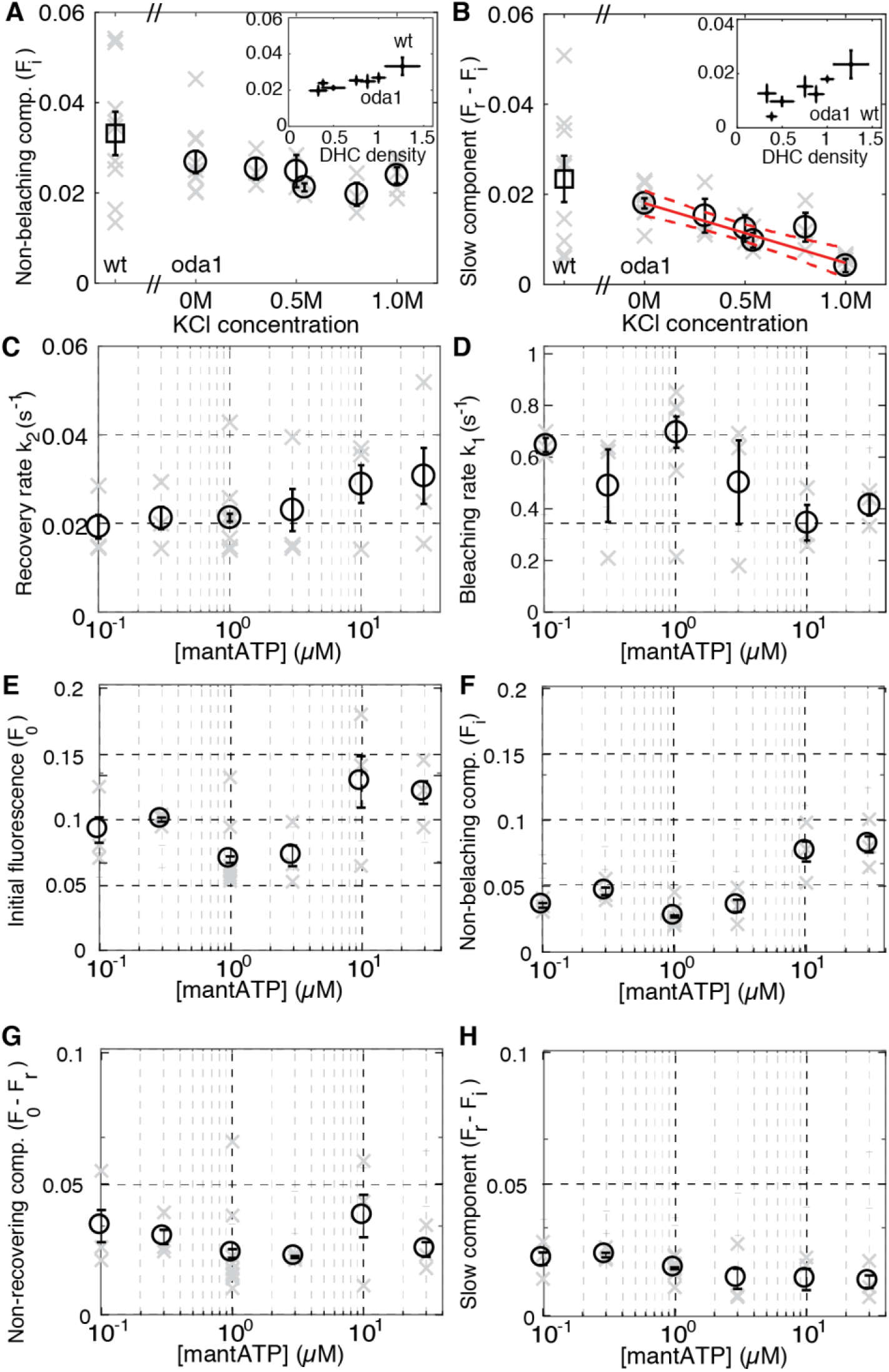
Dependencies of the fast and slow components on dynein content and mantATP concentration. **A** The non-bleaching component amplitude, *F_i_* at 1 μM mantATP. Inset shows the same data, plotted against measured density of DHC (using Figure 3). **B** Corresponding plot for the slow component amplitude, *F_r_ − F_i_* at 1 μM mantATP. Data plotted as mean ± standard error (left to right: *n* = 9, 10, 3, 2, 3, 3, 5). A linear regression through the salt data is shown in red with dashed lines indicating 95% confidence bounds. Inset shows same data, plotted against measured density of DHC. For both **A** and **B**, data are plotted as mean ± standard error (left to right: *n* = 9, 10, 3, 2, 3, 3, 5). The mean and SE for wild-type axonemes is indicated with a square marker to the left of the x-axis break. Amplitudes for the salt-extracted axonemes are plotted against the salt concentration to the left of the *x*-axis break. **C** Plot of recovery rate constant *k*_2_ against mantATP concentration on a semi-logarithmic scale. **D** Plot of bleaching constant *k*_1_ against mantATP concentration on a semi-logarithmic scale. **F** Intensity of non-bleaching component *F*_0_ plotted against mantATP concentration. **G** Intensity of non-recovering component *F*_0_ − *F_r_* plotted against mantATP concentration. **H** Relative intensity of slow component *F_r_ − F_i_* plotted against mantATP concentration. For **C-H**, data are shown as mean ± standard error (*n* = 3, 3, 10, 3, 3, 3).

Because the slow-recovering component was smaller in salt-extracted *oda1* axonemes than in non-extracted *oda1* axonemes, it is likely that mantATP is binding to inner-arm dyneins. It is also possible that mantATP also binds to outer arm dynein; however, the large variability of the rate for wild-type axonemes precluded us from making a definitive statement one way or the other.

### The bleaching and recovery rates as well as the amplitudes of the three components showed little dependence on mantATP concentration

To test whether the slow component is limited by the availability of mantATP, we varied the concentration of mantATP in the assay. Imaging conditions allowed us to vary the fluorescent nucleotide concentration only over a range of 0.1 μM to 30 μM. At the lowest concentration, the axonemes were so faint as to be barely visible against the background; at the highest concentration, the background was so bright that the axoneme could be barely discerned.

The recovery rate constant showed little dependence on the mantATP concentration (Figure 5C). This is expected because the unbinding of the bleached mantATP is not expected to be influenced by the mantATP in solution (Equation 12). The bleaching rate was also little affected by mantATP (Figure 5D). For the non-recovering site this is consistent with its very slow release of bleached mantATP (>100 s). For the slow-recovering site, the lack of dependence on mantATP implies that even at 0.1 μM most of the binding sites must be occupied (Equation 10), implying a high affinity *K*_D_ = *k*_+_/*k*_−_ ≤ 0.1 μM. These data suggest that both the non-recovering and slow-recovering sites are high affinity, consistent with the relative amplitudes of the respective components showing little dependence on mantATP concentration (Figure 4E,F). The relative amplitude of the non-bleaching component increases somewhat with mantATP (Figure 4G), as does its absolute amplitude; this might indicate that there is a low affinity non-bleaching component. In summary, the bleaching and recovery curves show little dependence on the mantATP concentration.

## Discussion

Using UV-TIRF, we visualized mantATP binding to the axoneme and found nucleotide-binding sites distributed approximately uniformly along the axoneme. The bleaching and recovery experiments indicate, that there are at least three components: non-recovering, non-bleaching and slow-recovering.

We suggest that the following interpretations of these components, which have approximately equal amplitudes.

i. The slow-recovering component is a mantATP binding site on axonemal dynein that hydrolyzes mantATP and releases the product mantADP only slowly, with rate *u* = 0.023 *s*^−1^. The bleaching rate and the amplitude of this component depend only weakly on the mantATP concentration, indicating that the rate of binding of mantATP is much greater than the bleaching rate at all mantATP concentrations.
ii. The non-recovering component could correspond to a high-affinity binding site that unbinds mantATP or mantADP very slowly (on the timescale of the experiment; *τ*_u_ > 300 s) so that once bleached, there is no measurable recovery. This explains the absence of a dependence of the bleaching rate and the amplitude of this component on the mantATP concentration.
iii. The non-bleaching component has two interpretations. It could correspond to a very long-lasting fluorescent state of mantATP or mantADP that is resistant to bleaching (*τ_b_* > 300 s). Alternatively, this component could correspond to a rapidly exchanging, non-hydrolyzing site with unbinding and binding rates fast compared to the sampling rate of 0.1 s.

If the slow-recovering component corresponds to axonemal dynein, and the slow recovery is due to the unbinding of mantADP, this could account for the slow beating of *Chlamydomonas* axonemes in mantATP, as previously proposed (23). The hydrophobic methylanthraniloyl substituent on the ribose may make the affinity for the analogue to the hydrophobic dynein binding pocket higher, reducing the off-rate. We found that isolated demembranated *oda1* axonemes do not beat at mantATP concentrations at or below 60 μM (24). At 100 μM mantATP they reactivate and have a beat frequency of 0.4 Hz (24). Note that the mantATP requirement for *Chlamdomonas* is much higher than previously reported for sea-urchin sperm, which has a beat frequency of 0.3 Hz at 2.5 μM mantATP (23). Unfortunately, we could not study the binding and bleaching dynamics of *Chlamydomonas* axonemes at high enough concentrations to reactivate the beat because of the poor signal-to-noise ratios. If one mantATP is hydrolyzed per dynein per beat cycle, at 100 μM mantATP we expect a dynein turnover rate of 0.4 s^−1^, and this must be faster than all rate constants between different states within the cycle, including mantADP unbinding. However, the recovery rate in our measurements was ~0.02 s^−1^. If this rate corresponds to mantADP release, then release would need to be accelerated several fold for beating at 0.4 Hz to occur. Therefore, we propose (assuming that our measured recovery rate corresponds to mantADP release) that either beating itself accelerates mantADP release, or that preventing beating through immobilization on the surface inhibits mantADP release. Thus our observations are consistent with axonemal dyneins showing force-dependent inhibition, as has been proposed by (12).

Directly observing labeled nucleotides bound to the axoneme is a promising method for characterizing dynein activity in the intact axoneme. If the fluorescence could be resolved in space and time, the traveling wave of activation of dynein could, in principle, be measured. The biggest difficulty currently is that dynein is very selective for which nucleotides it can utilize. We found that mantATP gives very low signals. Development of new fluorophores and conjugation methods is ongoing as demand grows for new and more stable fluorescent dyes. This gives us is hope that a fluorescently labeled ATP suitable for dynein, with better optical characteristics, may be available in the future.

The use of TIRF microscopy assisted us in reducing the background of diffusing labeled ATP in solution by illuminating only about 200 nm from the surface of the cover glass. The method works very well for observing axonemes that are attached to the surface. Once the axoneme is beating, it will likely be too far from the surface of the glass chamber for this method. It must also be noted that the beat of the axoneme is not planar. It is more suitable to use epifluorescent illumination for observations of fluorescence in the beating axoneme. To minimize background noise from diffusing fluorophore, a spiking assay, using a mixture of labeled and unlabeled ATP could be used. This will require using a fluorescent dye that allows near single-molecule dye resolution.

There are other important aspects to consider if one is to perform spatio-temporally resolved observation of dynein activity in a beating axoneme. The dynein proteins may be active all the time. In the case that the motors are at least down-regulated when not producing force, there would still be a possibility of resolving the wave of force. The motors are also likely to be alternately active on different sides of the axoneme, making it harder to resolve their activity.

Careful separation of contributions from dynein and other proteins is also important, as well as being aware of the existence of regulatory sites. Further studies on mutant axonemes with various non-dynein components missing might shed light on the source of the non-dynein components and possible kinase or non-dynein motor activity in the axoneme. We hope that with the current investigation we lay the groundwork for future successful observation of dynein in the intact beating axoneme.

## Appendix

In this section, we solve the kinetic scheme shown in Figure 2 under simplifying assumptions. The total concentration of binding states is

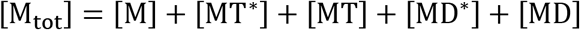

where M corresponds to the nucleotide binding protein in the apo state, T* is fluorescent mantATP, D* is fluorescent mantADP, T is bleached mantATP and D is bleached mantADP. *k*_+_ is the second-order rate constant for mantATP binding to M, and *k*_−_ is the unbinding rate. For *Tetrahymena* outer-arm dynein, Holzbaur & Johnson (1989) found *k*_+,ATP_ = 4 μM^−1^s^−1^ and *k*_−,ATP_ = 0.15 s^−1^ for ATP binding and unbinding respectively. We assume that [T*] is constant since the TIRF illumination depth (~0.2 μm) is only a small fraction of the 100-μm depth of the flow cell, and so bleaching of the mantATP in solution will be ≈500 times slower than that in the TIRF field. As a shorthand, we write *k* = *k*_+_[T*], which is 4 s^−1^ at 1 μM ATP. The rate constant h combines hydrolysis and phosphate (P_i_) release and is irreversible because the phosphate concentration in solution is negligible. For the axonemal dynein ATPase, hydrolysis is fast and reversible (rates of 100 s^−1^ and 30 s^−1^ respectively), and phosphate release is also fast (80 s^−1^) (Holzbaur & Johnson,1989): we therefore denote MT* and MT as combined ATP and ADP·P_i_ states with 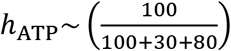 80 s^−1^ ≈ 40 s^−1^. *u* is the unbinding rate of the mantADP (D* and D), assumed not to depend on whether mantADP is bleached or not; the reverse rate is zero because the concentrations of D^*^ and D in solution are negligible. *u*_ADP_ ≈ 4 s^−1^ is the rate limiting step for the dynein ATPase, and is increased up to 5-fold in the presence of microtubules (Holzbaur & Johnson, 1989). *b* is the bleaching rate, which is irreversible.

We now make the simplifying assumption that *h* ≫ *k*_−_,*b,u*. This assumption is true for ATP (given that the bleaching rate is *b* ≈ 0.6 s^−1^, Table 2), and will be true for mantATP provided that hydrolysis is not disproportionally decreased. With this assumption, the kinetic scheme reduces to:

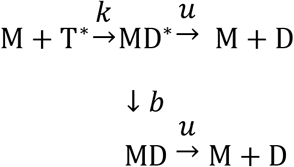

This has a simple solution. The steady-state fluorescence in the absence of bleaching is

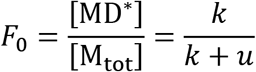

and in the presence of bleaching

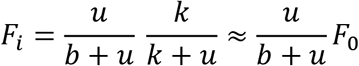

There are two rate constants, *k + u* and *b + u*: taking *k* large, we obtain the rate constants *k*_1_ = *b + u* in the presence of bleaching and *k*_2_ = *u* in the absence of bleaching (i.e. during recovery). Thus,

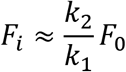

as stated in the main text following Figure 2.

## Author contributions

MF and JH designed the experiments. MF preformed the experiments and image and data analysis. JH and MF constructed the model. MM designed and MM and MF built the microscope setup. MF and JH wrote the manuscript in consultation with MM.

## Acknowledgements

The authors would like to thank Dr. Mark Kittisopikul for useful suggestions about the image analysis, and Dr. Veikko Geyer for discussions and advice throughout the course of this work.

## Abbreviations

ADP –: adenosine diphosphate
ATP –: adenosine triphosphate
MantATP -: methylanthraniloyl adenosine triphosphate
TIRF –: total-internal-reflection fluorescence

